# Cochlear implant material effects on inflammatory cell function and foreign body response

**DOI:** 10.1101/2022.06.20.496419

**Authors:** Megan J. Jensen, Alexander D. Claussen, Timon Higgins, Rene Vielman-Quevedo, Brian Mostaert, Linjing Xu, Jonathon Kirk, Marlan R. Hansen

## Abstract

**Objectives:** The objectives of this study were to assess the effects of cochlear implant (CI) biomaterials on the function of macrophages and fibroblasts, two key mediators of the foreign body response (FBR) and to determine how these materials influence fibrous tissue growth and new bone formation within the cochlea.

**Methods:** Macrophages and fibroblasts were cultured on polydimethylsiloxane (PDMS) and platinum substrates and human CI electrodes *in vitro*. Cell count, cell proliferation, cytokine production, and cell adhesion were measured. CI electrodes were implanted into murine cochleae for one week without electrical stimulation. Implanted cochleae were harvested for 3D X-ray microscopy with the CI left *in-situ*. The location of new bone growth within the scala tympani (ST) with reference to different portions of the implant (PDMS vs platinum) was quantified.

**Results:** Cell counts of macrophages and fibroblasts were significantly higher on platinum substrates and platinum contacts of CI electrodes. Fibroblast proliferation was greater on platinum relative to PDMS, and cells grown on platinum formed more/larger focal adhesions. 3D x-ray microscopy showed neo-ossification in the peri-implant areas of the ST. Volumetric quantification of neo-ossification showed a trend toward greater bone formation adjacent to the platinum electrodes compared to areas opposite or away from the platinum electrode bearing surfaces.

**Conclusions:** Fibrotic reactions are biomaterial specific, as demonstrated by the differences in cell adhesion, proliferation, and fibrosis on platinum and PDMS. The inflammatory reaction to platinum contacts on CI electrodes likely contributes to fibrosis to a greater degree than PDMS, and platinum contacts may influence the deposition of new bone, as demonstrated in the *in vivo* data. This information can potentially be used to influence the design of future generations of neural prostheses.

## 1. Introduction

Cochlear implants (CIs) are neural prosthetic devices used to restore hearing to patients with severe to profound hearing impairment. As is the case with all other implanted prosthetics and medical devices, placement of a CI initiates the foreign body response (FBR), which can result in fibrous tissue and new bone development/ossification within the cochlea(Foggia et al., 2019; Seyyedi and Nadol, 2014). Intracochlear fibrosis hinders performance for CI users by increasing electrode impedances, reducing dynamic range of stimulation, and interfering with CI battery life(Ishai et al., 2017). It has also been linked to delayed hearing loss after CI(Quesnel et al., 2016; Scheperle et al., 2017). Therefore, the FBR and its sequalae represent an area of research that warrants continued efforts to overcome these limitations.

The FBR involves adsorption of blood proteins, such as fibrinogen, to the biomaterial surface, followed by infiltration and recruitment of macrophages and other inflammatory cells to the site of the implant(Anderson et al., 2008; Williams, 2009). Appropriate surface protein adsorption enables macrophages to adhere to the biomaterial surface and undergo cell-cell fusion to form foreign body giant cells (FBGC)(Anderson et al., 2008). Macrophages and FBGCs release soluble mediators, such as cytokines and chemokines, which recruit additional inflammatory cells, including fibroblasts(Anderson et al., 2008). Macrophages and fibroblasts collectively influence extracellular matrix (ECM) remodeling. In normal wound healing, ECM remodeling facilitates restoration of normal tissue or scar formation. However, in the presence of a foreign body, the myofibroblast positive feedback loop becomes dysregulated and excessive ECM is deposited, ultimately leading to fibrous encapsulation of the biomaterial(Witherel et al., 2019).

In the cochlea, the acute portion of the FBR is attributed to trauma associated with electrode insertion(Knoll et al., 2022; Li et al., 2007). Other factors, such as bone dust and blood contamination within the cochlea may also play a role(Clark et al., 1995). The delayed, chronic portion of the FBR, however, is thought to persist due in part to the indwelling foreign material within the cochlea. Cochlear implants are comprised of platinum-iridium contacts encased in a polydimethylsiloxane (PDMS) carrier, of which both materials have been indirectly shown to contribute to the FBR. Consistent with this hypothesis, platinum and PDMS particles have been found within macrophages and the extracellular space of the middle and inner ear of human cochlear implant recipients(Nadol et al., 2014; Okayasu et al., 2020).

Although the roles of macrophages and fibroblasts in the FBR have been defined, little is known regarding how these cells interact with specific CI biomaterials (PDMS, platinum) and how the physical and chemical properties of CI biomaterials modulate the intensity of the FBR. Further insight into CI biomaterial mediated fibrosis is warranted and has the potential to assist in overcoming the current limitations of auditory prostheses. For example, thin film coatings could be applied to the surface of specific CI biomaterials to alter the surface properties and limit the FBR(Leigh et al., 2017; Leigh et al., 2019). The objective of this study was to assess the effect of CI biomaterials on the function (e.g., adhesion, proliferation, cytokine expression) of macrophages and fibroblasts, two key mediators of the FBR, and to determine how these materials influence fibrous tissue growth and new bone formation within the cochlea.

## 2. Materials and Methods

### 2.1. Materials

Polydimethylsiloxane (PDMS) kit Sylgard 184 was obtained from Dow Corning. Medical grade PDMS was obtained from Bentec Medical. Platinum-Iridium (80%/20%) sheets were obtained from Surepure Chemetals. Paraformaldehyde, collagenase, Dulbecco’s modified Eagle’s medium (DMEM), DMEM/F12, phosphate-buffered saline (PBS), and L-glutamine were obtained from Sigma-Aldrich. Recombinant Anti-vimentin antibody, Anti-F4/80 antibody, anti-phosphoFAK antibody, and Total Collagen Assay Kit were obtained from Abcam. DAPI-containing mounting medium, recombinant human granulocyte/macrophage colony-stimulating factor (GM-CSF), fetal bovine serum (FBS) and TrypLE Express were obtained from Gibco. 488 Goat anti-Rabbit secondary antibody and 546 Goat anti-Rat secondary antibody, Click-iT EdU Kit 647, and glass cover slips were obtained from ThermoFisher. The Mouse Cytokine Array Kit was from R&D Systems. The Cochlear Practice Electrodes (Slim Straight electrodes) were provided by Cochlear Corp.

### 2.2 Cell count on PDMS and Platinum

Macrophages and fibroblasts were cultured on PDMS and platinum substrates (7 x 7 x 0.15 mm) to determine differences in cell count. The substrates were disinfected with 70% ethanol, rinsed, and sterilized using UV radiation for 30 minutes prior to use. Bone marrow derived macrophages were obtained from 4-week-old CBA/J mice, as previously described.(Trouplin et al., 2013) Use of vertebrate animals was approved by and performed in accordance with the Institutional Animal Care and Use Committee at the University of Iowa (Iowa City, IA). The macrophages were maintained in macrophage complete medium (DMEM/F12 with 10% FBS, 10mM L-glutamine, 100 units/mL GM-CSF). Macrophages were seeded onto PDMS and platinum substrates at a density of 1.2 x 10^6^ cells/mL. The cells were cultured for 7 days, followed by fixation with 4% PFA diluted in PBS for 20 minutes at 4°C. They were then permeabilized with blocking buffer (1% BSA, 0.3% Triton-X) for 30 minutes at room temperature, followed by incubation with F4/80 antibody (Abcam, 1:100) for 2 hours at 37 °C. Secondary antibody (ThermoFisher Alexa-Fluor 488, 1:400) was then applied for 1 hour at room temperature. The substrates were cover slipped with DAPI containing mounting medium. Ten random 20x images of separate areas were taken per substrate using an epifluorescent microscope. Cells were counted manually. Three replicates were performed for each condition.

Cochlear fibroblasts were dissected from the spiral ligament of perinatal (p2-5) mouse pups. In detail, the spiral ligament was removed and digested in 1:1 trypsin and collagenase at 37°C for 20 minutes, with intermittent shaking. FBS was used to terminate the reaction. The cells were triturated, plated onto 35 mm culture dishes, and cultured in DMEM with 10% FBS until approximately 80% confluent. After 48-72 hours, the cells were trypsinized with TrypLE Express (ThermoFisher) for 10 minutes at 37° C. They were then centrifuged and plated onto PDMS and platinum substrates at a cell density of 2×10^4^ cells/mL. The fibroblasts were cultured for 48 hours, followed by fixation and immunostaining with anti-vimentin antibody (Abcam, 1:200) and secondary antibody (Thermofisher Alexa 488, 1:400). Nuclei were stained with DAPI containing mounting medium. Ten random 20x images were taken per substrate and cell count was determined, as before. Three replicates were performed for each condition.

### 2.3 Cell count on human CI electrodes

3T3 mouse embryonic fibroblast cell count was also measured on human CI electrodes (Cochlear Practice Electrode, Slim Straight electrodes) to determine if differences in substrate shape (cylindrical electrodes versus flat PDMS/platinum substrates) yielded similar results. The electrodes were divided into segments containing two platinum contacts and two equal length segments of PDMS. The electrodes were sterilized with 70% ethanol and 30 minutes of UV light. 3T3 fibroblast cultures were maintained for 48 hours, followed by immunostaining and determination of cell count, as before. CI electrodes were flattened, and z-stack was used to facilitate cell counting on curved areas of the implant. The area measured per electrode was 150 μm x 200 μm. Four replicates were performed for each condition.

### 2.4 Cell proliferation

To determine differences in macrophage and fibroblast proliferation on PDMS and platinum, macrophages and fibroblasts were cultured on PDMS and platinum substrates. After 7 days (macrophages) or 48 hours (fibroblasts), the cells were incubated in 5-Ethynyl-2’-deoxyuridine (EdU) for 2 hours. EdU was diluted (1:200) in the appropriate macrophage or fibroblast medium (macrophage complete medium and DMEM with 10% FBS, respectively). After incubation in EdU, the cells underwent EdU detection (ThermoFisher, Click-iT EdU Kit Alexa Fluor 647) followed by cell fixation, permeabilization, and immunostaining with anti-F4/80 antibody (1:100) and anti-vimentin (1:200) antibody for macrophages and fibroblasts, respectively. Ten randomly selected 20x images of separate areas were taken using an epifluorescent microscope. Cells were counted manually. Cell proliferation was measured by counting the number of EdU positive (Alexa Fluor 647 expressing nuclei) macrophages or fibroblasts relative to the total number of cells. Three replicates were performed for each condition.

### 2.5 Cytokine production and macrophage-fibroblast co-cultures

To further understand how CI biomaterial surfaces influence cell signaling, specifically cytokine/chemokine production, macrophages were cultured on PDMS and platinum substrates for 7 days. On day 7, half of the cultures underwent an endotoxin challenge with 25 ng/mL of lipopolysaccharide (LPS) for 8 hours to prime the cells.(Deng et al., 2013) LPS was selected due to its ability to stimulate macrophages to become classically activated.(Witherel et al., 2019) Medium from the cultures without LPS was used as a control. After 8 hours, 1.5 mL of medium from each condition was collected for cytokine analysis and relative intensity of a panel of cytokines was measured (R&D Systems, Mouse Cytokine Array Kit).

To further investigate the effect of CI biomaterials on macrophage signaling and the influence of macrophages on fibroblast proliferation, cochlear fibroblasts were added directly to established, mature macrophage cultures to create macrophage – fibroblast co-cultures. Fibroblast cultures (without macrophages) were used as controls. Co-cultures were maintained in macrophage complete medium for 48 hours, then incubated with EdU for 2 hours, as before. After EdU detection, co-cultured substrates were immunostained with anti-F4/80 antibody (1:100; macrophage labelling) and anti-vimentin antibody (1:200; fibroblast labelling), with 1:400 Alex Fluor 546 and 1:400 Alexa Fluor 488 secondary antibodies, respectively. The substrates then underwent EdU detection protocol and were cover slipped with DAPI containing mounting medium. Control substrates were immunostained with anti-vimentin antibody, secondary antibody (1:400 Alexa Fluor 488), and DAPI. Ten 20x images were taken per substrate. Fibroblasts were differentiated from macrophages by splitting the color channels in Image J (fibroblasts were identified using the green fluorescent protein (GFP) channel, macrophages were identified using the tetramethylrhodamine (TRITC) channel). Fibroblasts were counted manually. The percentage of EdU positive fibroblasts relative to total fibroblasts was measured. Three replicates were performed for each condition.

### 2.6 Cell adhesion

Differences in cell adhesion are based on the binding of proteins to biomaterial surfaces and thus are dependent on surface material properties (Witherel et al., 2019). One way to measure cell adhesion is through the phosphorylation of focal adhesion kinase (FAK). Therefore, anti-phosphorylated FAK expression was used to detect differences in focal adhesion dynamics. Macrophages and fibroblasts were cultured on PDMS and platinum for 4 hours. The cells were then fixed, permeabilized and immunostained with anti-phosphorylated focal adhesion kinase antibody (anti-phospho-FAK, ThermoFisher, 1:200) followed by secondary antibody (ThermoFisher, Alexa-Fluor 633, 1:400). Cell nuclei were stained with DAPI. Using epifluorescent microscopy, 30 randomly selected 63x images were taken per substrate. Using Image J software, the number and size of focal adhesions per cell was measured, as well as the average area occupied by the focal adhesions using a previously described method.(Horzum et al., 2014)

### 2.7 3-D X-ray microscopy of in-vivo mouse cochlear implants

We have previously described robust intracochlear fibrous tissue and new bone growth in mouse cochleae following CI placement(Claussen et al., 2019). In a separate set of *in-vivo* experiments, 3D X-ray microscopy was used to observe volumetric and spatial patterns of new bone growth influenced by the local proximity to different implant materials (PDMS carrier vs platinum contacts).

10-week old CBA/CaJ (n=6) mice were unilaterally (left ear) implanted through round window insertion with a custom 3 electrode cochlear implant (approximate insertion depth of 2.25mm) as previously described(Claussen et al., 2019). No electric stimulation was performed during the duration of implantation. Subjects were singly housed in the standard University of Iowa murine housing system for the duration of the experiments and given ad-libitum access to food and water. Following 21 days of implantation, subjects were sacrificed, and the left implanted cochleae harvested for 3D X-ray microscopy with the CI left *in-situ*. Cochleae were osmicated to enhance soft tissue contrast and not de-calcified to enhance bony detail for 3D X-ray microscopy as previously described(Claussen et al., 2019).

Cochleae were imaged using the Zeiss Xradia Versa 3D X-ray microscope (Zeiss, USA), producing a voxel size of 1.5-2μm. Initial image series were obtained with the CI *in-situ*, with another image series obtained immediately after CI removal. Subsequent image merging, 3-dimensional reconstruction, volumetric segmentation and analysis was performed in Dragonfly 4.1 (ORS, Canada). These image series were manually merged using durable bony landmarks to create a composite image projecting the CI within the cochlea without associated artifact. 3D volume segmentation of the implant, areas of bone and soft tissue growth within the scala tympani were performed by an experimenter blinded to the sample ID. The location of new bone growth within the scala tympani with reference to different portions of the implant and proximity to potential FBR inducing materials (PDMS vs platinum) was quantified within 3 separate periimplant locations: “electrode”; the 180 degree peri-implant area adjacent to the half banded platinum electrode surface in the electrode bearing portion of the CI, “anti-electrode”; the 180 degree peri-implant area adjacent to the PDMS only surface in the electrode bearing portion of the CI, and “non-electrode”; the 360 deRgree peri-implant area outside of the electrode bearing areas of the CI.

### 2.8 Statistical analysis

Unpaired, two-tailed t-tests were used to compare cell growth, proliferation, and focal adhesion number, size, and area on platinum and PDMS. Ordinary one-way ANOVAs with post-hoc tukey tests were used to compare percentage of EdU positive cells and cytokine expression levels on PDMS versus platinum. 3D X-ray microscopy data were analyzed in GraphPad Prism software (GraphPad Software, USA) via ordinary one-way ANOVA.

## 3. Results

### 3.1 Cell count

To assess cell growth on CI biomaterial surfaces, bone-marrow derived macrophages and cochlear fibroblasts were plated on PDMS or platinum surfaces. Fig. 1 demonstrates differences in macrophage and fibroblast counts on PDMS and platinum. There were significantly higher counts of macrophages (two-tailed t-test, p=0.002, df=4) and fibroblasts (two-tailed t-test, p<0.001, df=38) on platinum relative to PDMS, with a 4-to-5-fold increase in cell count on platinum substrates.

**Figure 1.**
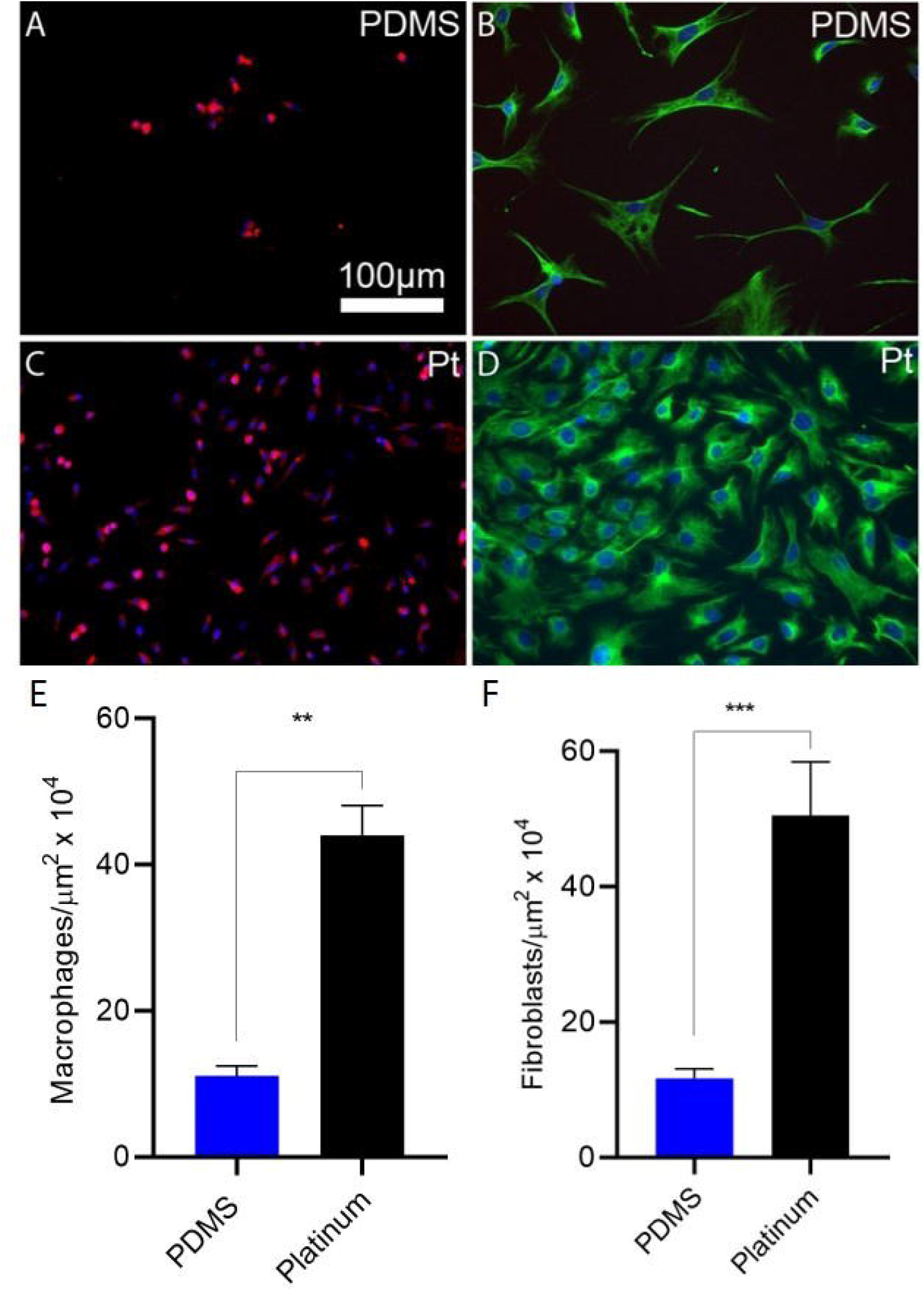
Cell growth on PDMS and platinum surfaces. Macrophages and cochlear fibroblasts were plated on PDMS and platinum and cell counts were measured after 7 days. Macrophages (A, C) were labeled with anti-F4/80 antibody (green) and DAPI (blue). Fibroblasts (B, D) were labeled with anti-vimentin antibody (green) and DAPI (blue). There were significantly fewer macrophages (E) and fibroblasts (F) on PDMS relative to platinum, p=0.002 and p<0.001, respectively. The experiment was repeated with at least 3 separate cultures. Error bars present standard error of the mean.

To further evaluate the adhesion and growth of cells on actual human CI electrode array biomaterials, we cultured 3T3 fibroblasts on human CI electrode array for 48 hours and quantified the density of cells on either the platinum electrode contacts or the adjacent PDMS housing (Fig. 2). A 2-to-3-fold increase in cell count was noted on the platinum contacts relative to the surrounding PDMS casing (p=0.0134).

**Figure 2.**
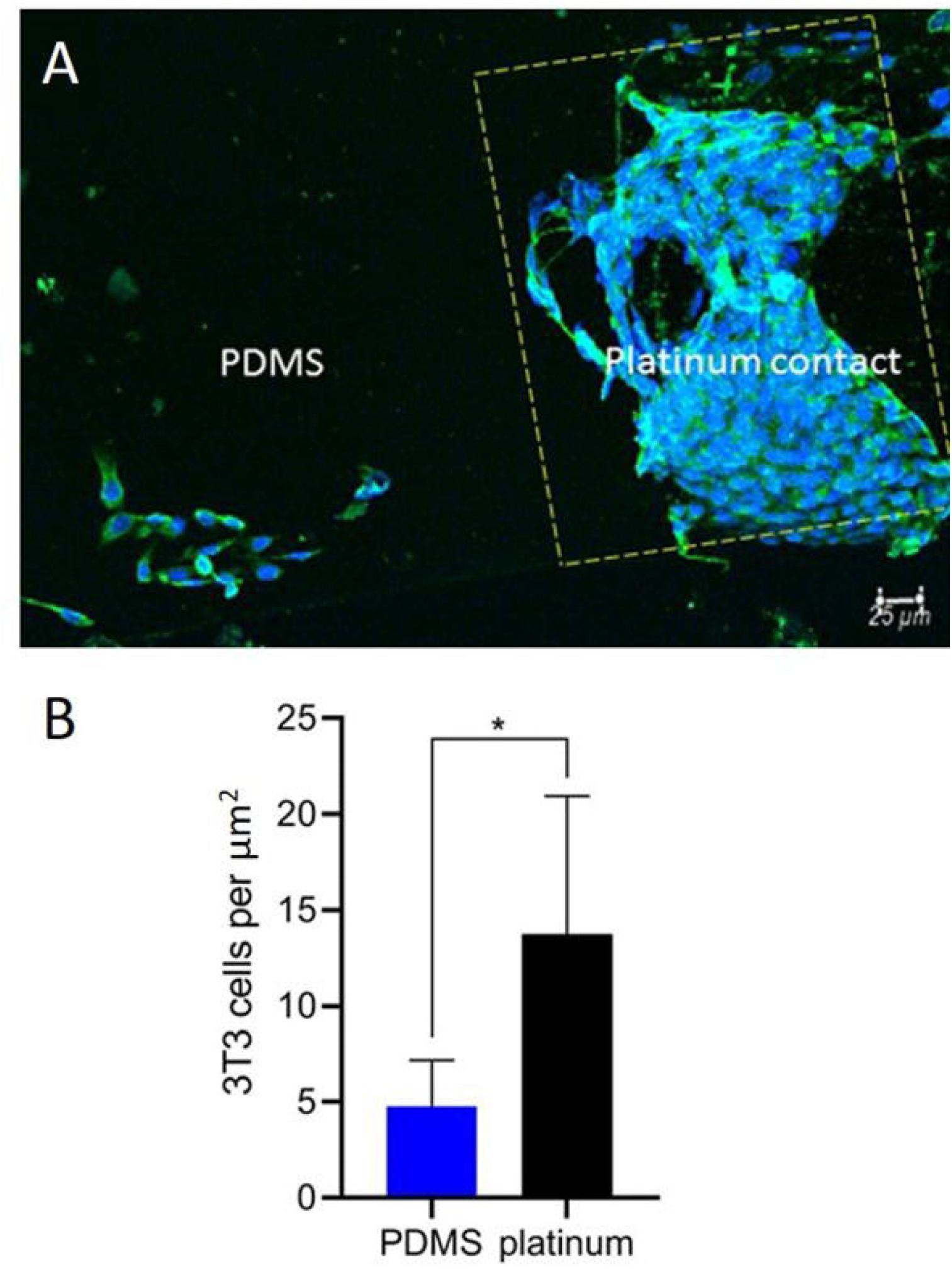
Cell count on human cochlear implant electrode array. 3T3 fibroblast cells were cultured on CI electrode arrays. 3T3 fibroblasts were labeled with anti-vimentin antibody (green) and DAPI (blue). The platinum contact is outlined by the yellow dashed line. There were significantly fewer 3T3 cells (B) on PDMS relative to platinum (p=0.0134, two tailed t-test). The experiment was repeated with four separate cultures. Error bars present standard error of the mean.

### 3.2 Cell proliferation

Differences in cell numbers after 7 days represents effects of both the number of initially adhered cells as well as the proliferation rate of the cell on each surface. Cell proliferation on the different surfaces was therefore directly assessed by determining the percent of cells that incorporate EdU, a thymidine analog, during the S phase of the cell cycle. Macrophage and fibroblast proliferation, measured as the percentage of EdU-expressing nuclei relative to total cells, is represented in Fig. 3. The percentage of EdU-expressing macrophage nuclei was similar on platinum and PDMS (two-tailed t-test, p=0.30, df=4). However, fibroblast proliferation differed significantly between the two materials, with a greater percentage of EdU-expressing fibroblasts in the platinum group relative to PDMS (two-tailed t-test, p=0.030, df=3).

**Figure 3.**
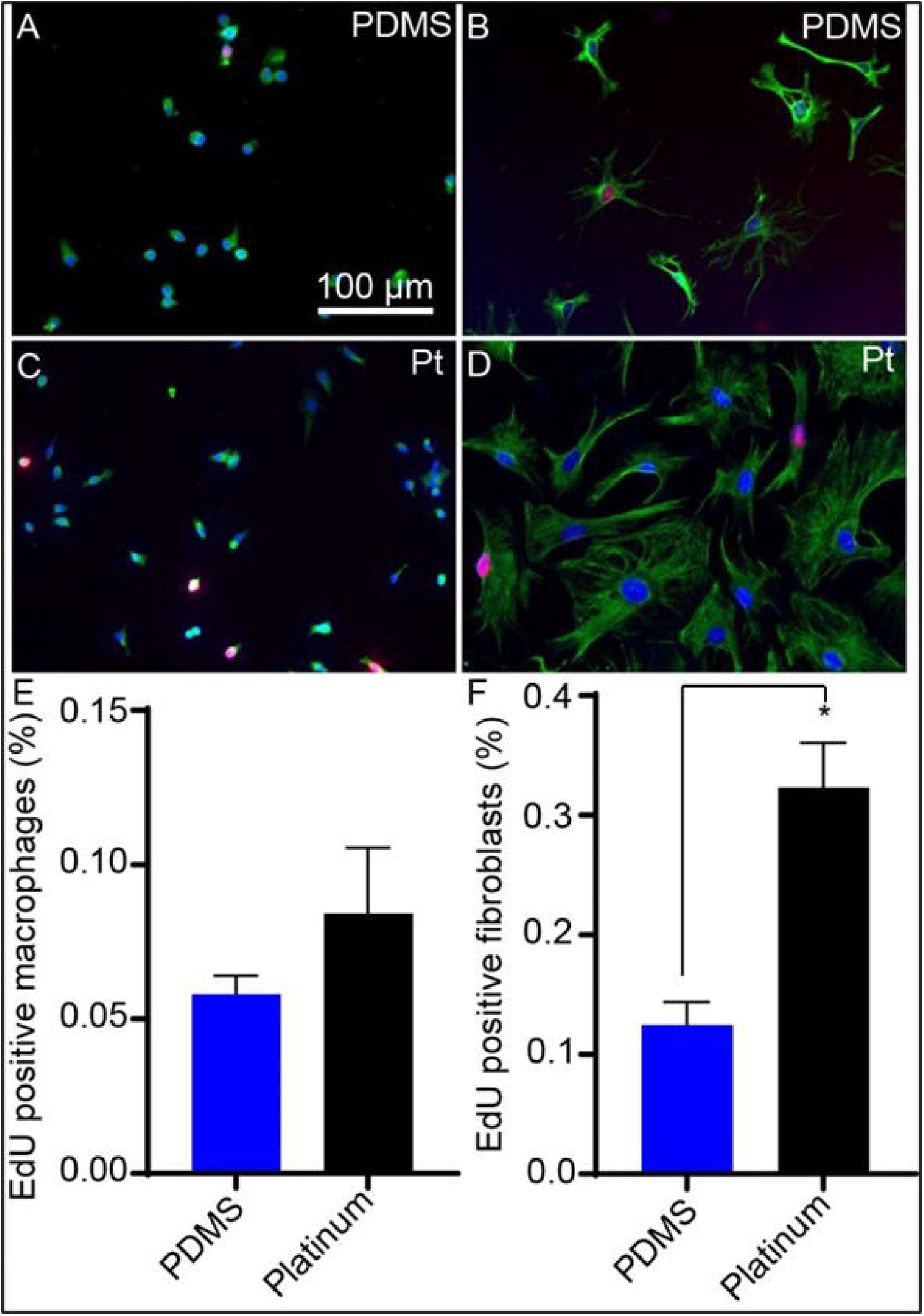
Cell proliferation on PDMS and platinum surfaces. Macrophages and fibroblasts were cultured on PDMS and platinum. Macrophages (A, C) were labeled with anti-F4/80 antibody (green) and DAPI (blue). Fibroblasts (B, D) were labeled with anti-vimentin antibody (green) and DAPI (blue). Cell proliferation was measured as the percentage of EdU-expressing (red) nuclei. Macrophage proliferation was similar on PDMS and platinum surfaces (E), p=0.30. Fibroblast proliferation differed significantly, with a greater percentage of EdU-expressing fibroblasts in the platinum condition relative to PDMS (F), p=0.030. The experiment was repeated with at least 3 separate cultures. Error bars present standard error of the mean.

### 3.3 Cell adhesion

To determine differences in the ability of cells to adhere to PDMS and platinum surfaces, active focal adhesions were assessed by labeling with anti-phosphorylated FAK antibodies and determining the number, size, and overall area of focal adhesion complexes for both macrophages and fibroblasts (Fig. 4). Macrophages formed a greater number of focal adhesions on platinum surfaces relative to PDMS (two-tailed t-test, p=0.031, df=12). The focal adhesions on platinum were larger on average (two-tailed t-test, p=0.003, df=12) and occupied a greater surface area (two-tailed t-test, p=0.0004, df=12) than those formed on PDMS. Similarly, fibroblasts formed a greater number of focal adhesions on platinum relative to PDMS (two-tailed t test, p=0.004, df=4). The focal adhesions occupied a greater surface area on platinum (twotailed t test, p=0.023, df=4), however there were no differences in the mean size of the focal adhesions between fibroblasts grown on PDMS and platinum (two-tailed t test, p=0.24, df=4).

**Figure 4.**
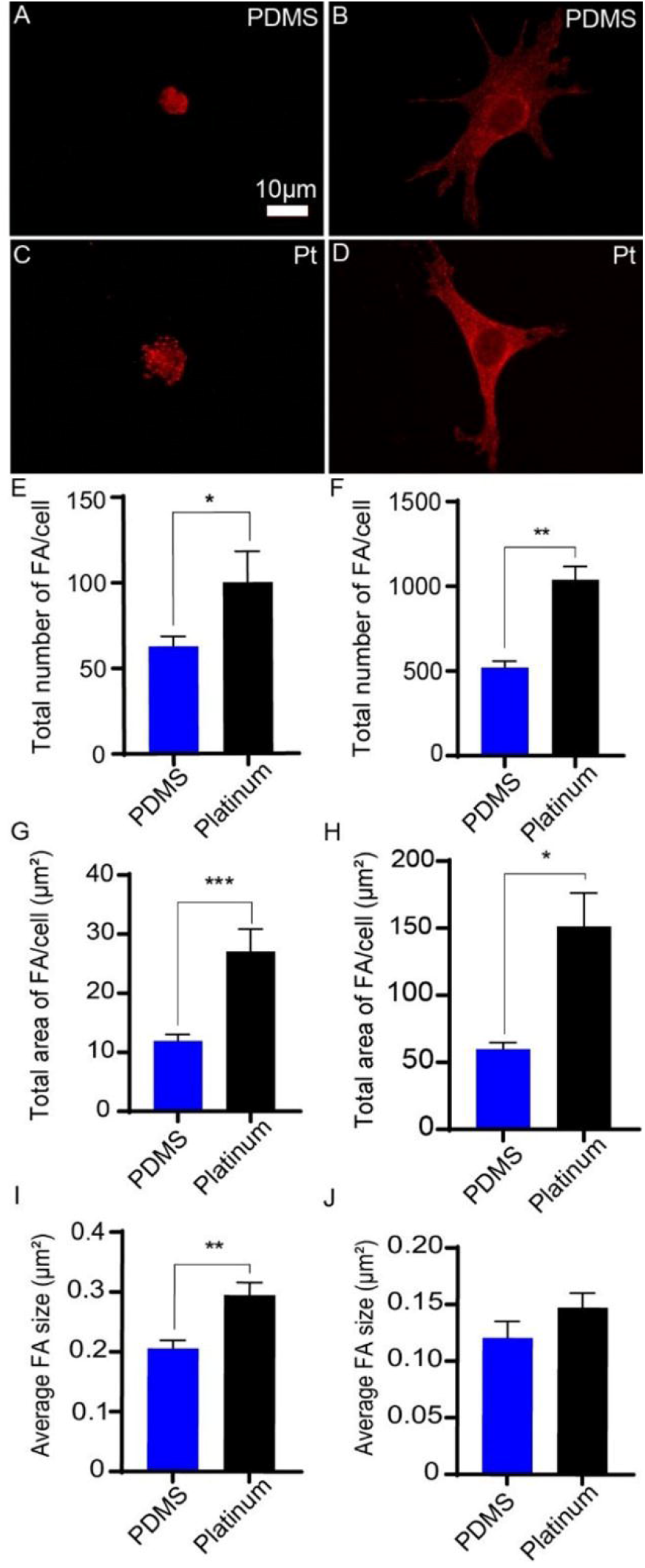
Focal adhesion formation on PDMS and platinum surfaces. Macrophages (A, C) and cochlear fibroblasts (B, D) were cultured on PDMS and platinum for 4 hours, followed by labeling with anti-phosphorylated FAK antibody (red). Macrophages formed a greater number of focal adhesions on platinum surfaces relative to PDMS (E), p=0.031, and the focal adhesions were larger on average (I), p=0.003, and occupied a greater surface area on platinum than on PDMS (G), p=0.0004. Similarly, fibroblasts formed a greater number of focal adhesions on platinum relative to PDMS (F), p=0.004. The focal adhesions occupied a greater surface area on platinum (H), p=0.023, however there were no differences in the mean size of the focal adhesions between fibroblasts grown on PDMS and platinum (J), p=0.24. The experiment was repeated with at least 3 separate cultures. Error bars present standard error.

### 3.4 Cytokine production and macrophage fibroblast co-cultures

Levels of chemokine/cytokine, interleukin, and pro-inflammatory marker expression by macrophages grown on PDMS and platinum with and without LPS priming are represented as relative intensities in Fig. 5. In the PDMS condition, increased levels of specific cytokines/chemokines, including CCL3 (p=0.019), CCL4 (p=0.001), CCL9 (p=0.007), CCL10 (p<0.0001), and CXCL2 (p=0.0001) were expressed when the macrophages were primed with LPS relative to non-LPS conditions. Expression levels of interleukins and other pro-inflammatory markers (e.g., C5/C5a, TNF-α, IFN-gamma, CD54, and TIMP-1) were similar between LPS and non-LPS groups (p>0.999) Cytokine/chemokine, interleukin, and other pro-inflammatory marker expression from macrophages grown on platinum did not differ between LPS and non-LPS groups (p>0.999). When comparing PDMS and platinum groups in the non-LPS condition, CXCL10 (p=0.005) and CCL3 (p=0.038) were the only chemokines/cytokines with increased expression in the platinum condition relative to PDMS. When comparing PDMS and platinum groups in the LPS-treated groups, IL-1ra was the only interleukin with significantly higher expression in the PDMS condition relative to platinum (p=0.044).

**Figure 5.**
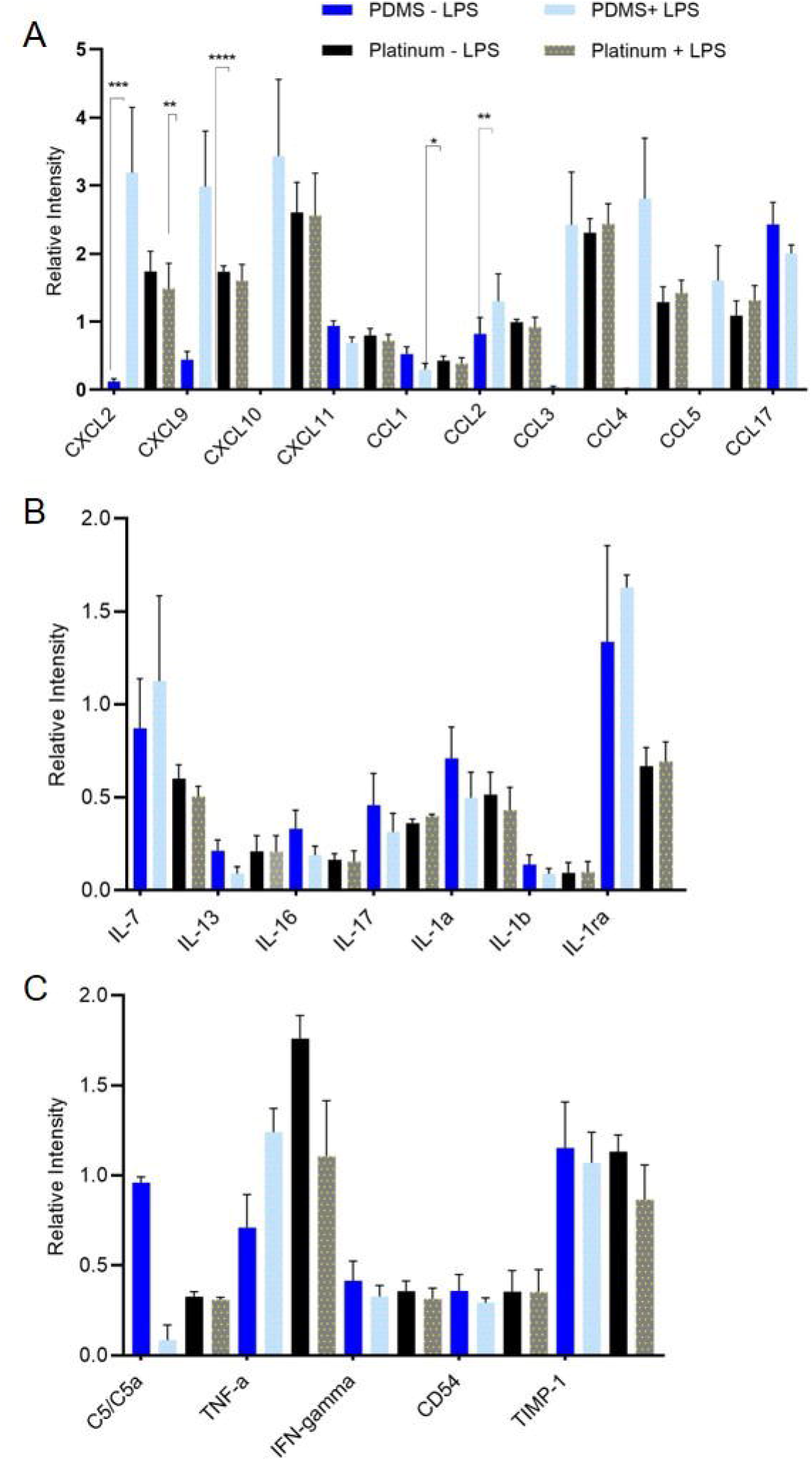
Cytokine expression on PDMS and platinum surfaces. Macrophages were cultured on PDMS and platinum for 7 days, then underwent an endotoxin challenge with the addition of 25 ng/mL of lipopolysaccharide (LPS) for 8 hrs to prime the cells. Medium without LPS was used as a control. After 8 hours, 1.5 mL of medium from each condition was collected for cytokine analysis. Chemokine/cytokine (A), interleukin (B), and other pro-inflammatory marker (C) expression is represented as a relative intensity. In the PDMS condition, increased levels of CCL3 (p=0.019), CCL4 (p=0.001), CXCL9 (p=0.007), CXCL10 (p<0.0001), and CXCL2 (p=0.0001) were expressed when the macrophages were primed with LPS. Expression levels of interleukins and other pro-inflammatory markers were similar between LPS and non-LPS groups. Cytokine/chemokine, interleukin, and other pro-inflammatory marker expression from macrophages grown on platinum did not differ between LPS and non-LPS groups. When comparing PDMS and platinum groups without LPS, CXCL10 (p=0.005) and CCL3 (p=0.038) were the only chemokines/cytokines with increased expression in the platinum conditions relative to PDMS. When comparing PDMS and platinum groups with LPS, IL-1ra was the only interleukin with significantly higher expression on PDMS relative to platinum (p=0.044). The experiment was repeated with at least 3 separate cultures. Error bars present standard error.

### 3.5 Effect of macrophages on fibroblast proliferation

The FBR involves a milieu of cells, predominantly macrophages and fibroblasts that interact and influence each other. To assess the effects of PDMS and platinum surfaces on macrophage stimulation of fibroblast proliferation, EdU uptake by fibroblasts on either PDMS or platinum was compared in co-cultures of macrophages and fibroblasts and also compared to fibroblast only cultures that lacked macrophages. In a subset of cultures, LPS was added to further stimulate the macrophages. Results from the macrophage-fibroblast co-cultures can be found in Fig. 6. On the PDMS surfaces, the addition of macrophages to create macrophage-fibroblast cocultures significantly increased fibroblast proliferation relative to the fibroblast-only condition. Both co-culture conditions, the LPS-treated condition (p=0.005) and non-LPS condition (p=0.043), had significantly more EdU expressing cells than the fibroblast-only controls. There was a trend toward increased EdU expression in the PDMS + LPS group relative to the PDMS non-LPS group, however this trend was not significant (p=0.098). On platinum, there were no significant differences in fibroblast proliferation between + LPS, non-LPS and control conditions (p=0.38).

**Figure 6.**
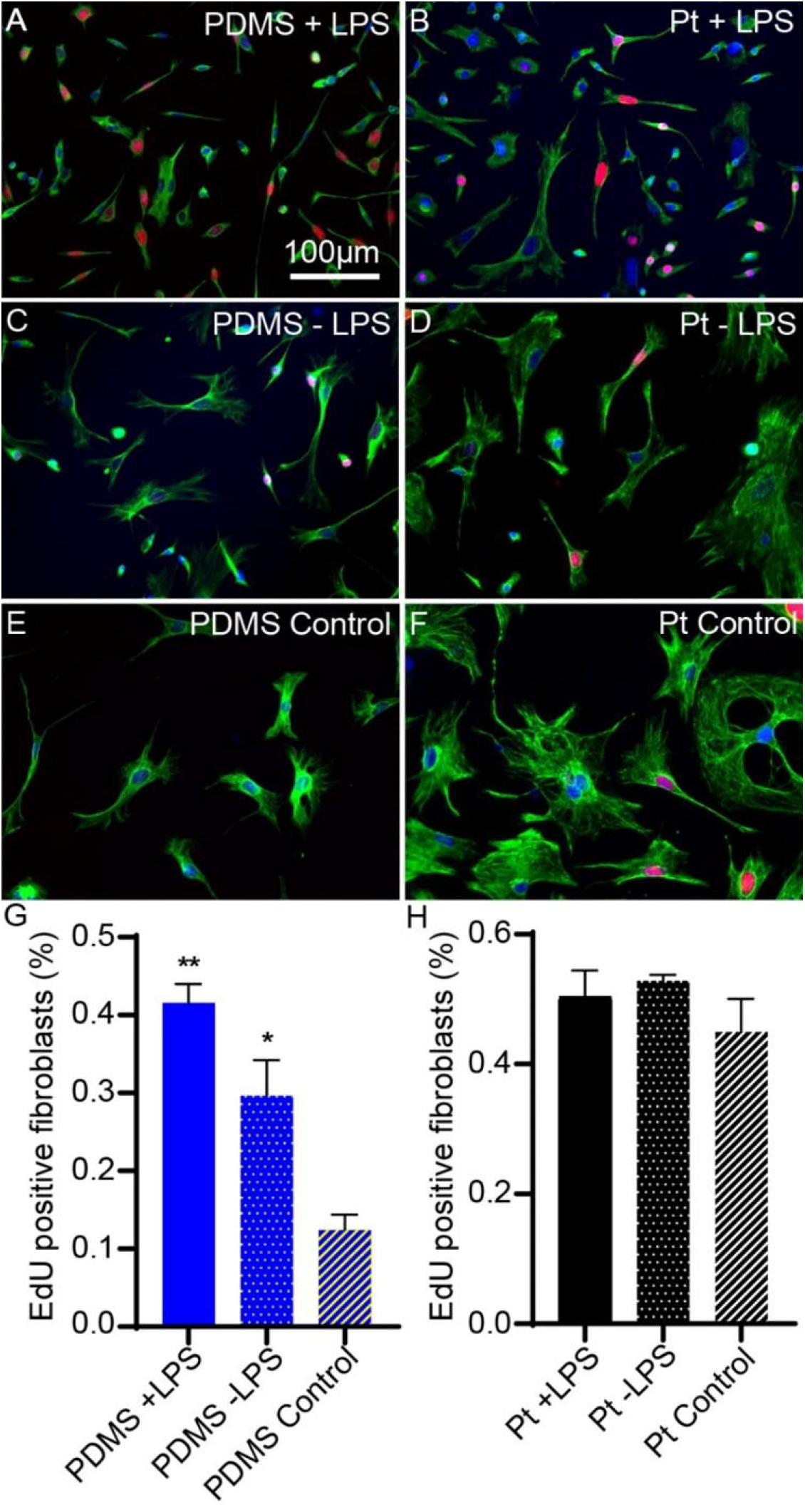
Effect of macrophages on fibroblast proliferation on PDMS and platinum surfaces. Macrophages and fibroblasts were co-cultured or fibroblasts were cultured in the absence of macrophages (control). Cell proliferation was measured as the percentage of fibroblasts with EdU-expressing (red) nuclei. Macrophages were labeled with anti-F4/80 antibody (not shown in these images) and fibroblasts were labeled with anti-vimentin antibody (green). Nuclei were stained with DAPI (blue). In the PDMS condition, macrophage-fibroblast co-cultures, either treated or not treated with LPS demonstrated increased fibroblast proliferation relative to the fibroblast-only control condition) (G) p=0.005 and p=0.043, respectively. There was a trend toward increased EdU incorporation in the PDMS + LPS group relative to the PDMS – LPS group, however this trend did not reach statistical significance, p=0.098. In the platinum condition, there were no significant differences in fibroblast proliferation between macrophage-fibroblast co-cultures + LPS, - LPS and control conditions (H), p=0.38. The experiment was repeated with at least 3 separate cultures. Error bars present standard error.

### 3.6 Intracochlear neo-ossification relative to PDMS and platinum surfaces

Neo-ossification represents the most advanced form of the FBR and is commonly seen in the scala tympani following CIs in humans and animal models. Here 3D x-ray microscopy was used to determine the location of intracochlear ossification relative to the PMDS and platinum surfaces of an implanted electrode array in mouse cochleae. All cochleae showed robust neoossification confined to the peri-implant areas of the scala tympani, with no bone formation seen distal to the implant tip or in other scala. A typical response is shown in Fig. 7, with most of the neo-ossification located adjacent to the implant as opposed to extending directly from the scalar walls. Volumetric quantification of neo-ossification showed a non-significant (p=0.5372, df=2) trend toward greater bone formation adjacent to the platinum electrodes (“electrode” 5.2 x 10^6^ μm^3^) compared to areas opposite (“anti-electrode” 3.8 x 10^6^ μm^3^) or away (“non-electrode 2.7 x 10^6^ μm^3^) from the platinum electrode bearing surfaces of the CI (Fig. 7). A more extensive soft tissue fibrotic response was seen in all cases within the scala tympani, extending up to the depth of CI insertion with an average volume of 8.1 x 10^7^ μm^3^. No evidence of traumatic CI insertion, including basilar membrane translocation or tenting or osseous spiral lamina fracture were seen in any subject.

**Figure 7.**
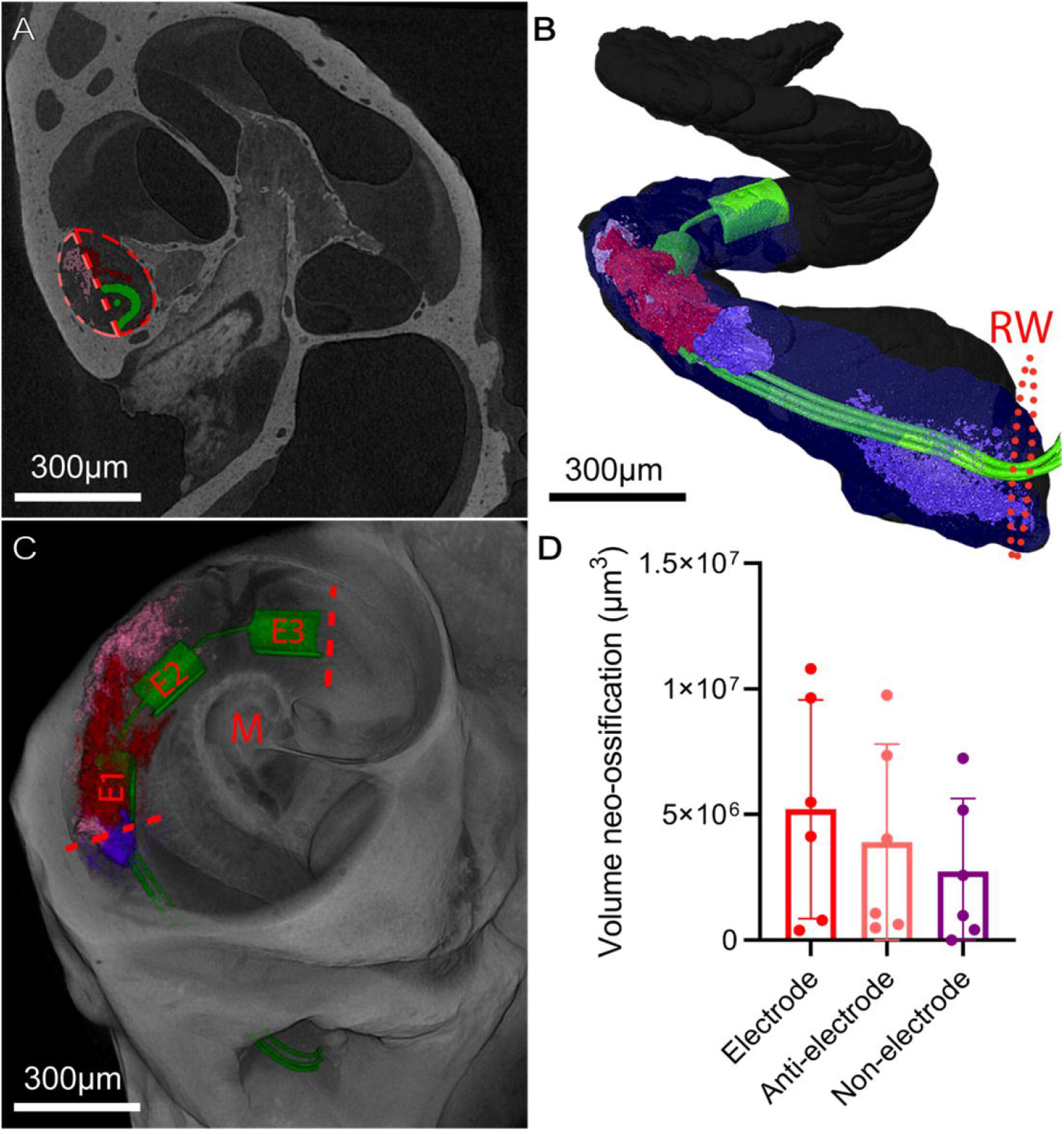
Intracochlear neo-ossification relative to platinum and PDMS bearing surfaces. 3D X-ray microscopy images of chronically implanted mouse cochleae (A,B,C) with the CI imaged *in-situ* were volumetrically segmented to compare the amount of neo-ossification in 3 separate areas: The areas immediately facing platinum electrode surfaces “Electrode” (red), PDMS surfaces opposing the electrode surface “Anti-electrode” (pink), and the intracochlear portions of the CI proximal to the most proximal E1 electrode “Non-electrode” (purple). The gray and blue shading in (B) represent the scala tympani space and areas of soft tissue fibrosis without neoossification, respectively.” Dashed lines in (A) denote the manual separation of the “Electrode” (red) and “Anti-electrode” (pink) areas in which neo-ossification was volumetrically quantified. The dashed line in (B) indicates the threshold of the round window. Dashed lines in (C) indicate the proximal and distal limits confining the “Electrode” and “Anti-electrode” areas, with the “Non-electrode” area being proximal to this area, stopping at the round window. The greatest volume of neo-ossification was seen in the “Electrode” area immediately adjacent to the Platinum electrodes, followed by the “Anti-electrode” and “Non-electrode” areas, however these differences were not significant (p>0.05).

## 4. Discussion

The insertion of a CI electrode array into the cochlea results in a FBR that can limit success for CI users(Quesnel et al., 2016). This response, comprised of macrophages, foreign body giant cells, fibroblasts, and other immune cells, ultimately leads to fibrosis and new bone formation within the scala tympani(Nadol et al., 2014). CI electrodes are comprised of platinum contacts encased in PDMS. It is felt that these materials collectively contribute to the inflammatory reaction within the cochlea(Nadol et al., 2014). However, differences in the cellular response to specific CI biomaterials have not been elucidated. The findings of this study further define the influence of PDMS and platinum on the FBR within the cochlea. Taken together, the results suggest that, compared to PDMS surfaces, platinum surfaces allow for greater cell attachment, growth, proliferation, and neo-ossification.

Our *in vitro* work demonstrated that cell growth, proliferation, adhesion, and cytokine synthesis are all factors affected by differences in biomaterial surface properties. For example, fibroblast counts were significantly greater on the platinum surfaces of CI electrode arrays, with far fewer cells adhering to the surrounding PDMS casing. While differences in geometry (shape/contour) of the platinum and PDMS surfaces of the CI electrode arrays may account for some differences in patterns of cell growth, it seems most likely that surface properties represent the driving force when viewed in combination with the cell behavior on flat surfaces.

When exposed to an opportune biomaterial surface, such as platinum, which is chemically inert/highly hydrophilic, fibroblasts will adhere and proliferate. PDMS, which is deemed chemically inert, has a low surface energy, and is relatively hydrophobic is often used in biomaterial applications either as a housing material as in CIs or as an implanted bulk material. The data in this study suggest that, compared to platinum, PDMS is a less favorable surface for cell growth and proliferation. However, based on our results, cells adhere and proliferate under the right conditions (presence of macrophages, pro-inflammatory signals, paracrine signaling, etc.) on PDMS, albeit to a lesser degree than that observed on platinum.

Cellular adhesion to a biomaterial surface is a complex process determined by underlying material/surface properties including stiffness, hydrophilicity, surface energy, and surface roughness, among other features(B. Majhy, 2021; Hallab et al., 2001; Razafiarison et al., 2018). Surface energy is generally significantly higher for metals compared to polymeric surfaces including PDMS, and cellular adhesion on metals demonstrates a linear correlation with surface energy(Hallab et al., 2001). Methods to alter surface energy or surface roughness have been explored in an effort to alter cellular responses to implanted biomaterials(B. Majhy, 2021; Hallab et al., 2001; Leigh et al., 2017; Leigh et al., 2019; Razafiarison et al., 2018).

Similar to the findings from the *in vitro* experiments, the *in vivo experiments* involving implanted mouse cochleae also demonstrated a trend toward greater neo-ossification in areas opposing the platinum surfaces of the array as opposed to PDMS surfaces. The soft tissue fibrotic response was seen to completely fill the scala tympani to the depth of the electrode tip, with no areas absent of fibrosis to allow a similar comparison between platinum and PDMS surfaces. Prior human and animal studies have demonstrated a similar trend toward greater tissue response around the platinum surfaces of the CI, however a key difference is that these prior studies included electric stimulation via the CI(Ishai et al., 2017; Shepherd et al., 2020). Electric stimulation was not included in the present study as we aimed to purely study any potential foreign body effects of the CI materials. Recent literature has demonstrated the role electric stimulation could play in exacerbating a platinum-related FBR secondary to irreversible dissolution of platinum microaggregates and platinum ions at high charge densities(Shepherd et al., 2020). Interestingly, islands of neo-ossification within the scala tympani, absent from the scalar walls were commonly seen, suggesting the process of de-novo neo-ossification within the scala as opposed to extension from the scalar endosteum (presumably from insertion trauma)(Shepherd et al., 2020). This finding of de-novo neo-ossification within the scala is not surprising as an influx of mesenchymal progenitors with chondrogenic potential associated with the inflammatory cell response in addition to the relatively hypoxic status of the scalar fluids could represent two factors predisposing to neo-ossification or “heterotopic ossification,” as has been documented within other tissues(Kan et al., 2018). Of note, other factors such as contamination with blood or bone dust(Clark et al., 1995), as well as damage to the lateral wall or osseous spiral lamina of the cochlea during insertion may contribute to initiation of neoossification(Kamakura and Nadol, 2016; Knoll et al., 2022; Li et al., 2007).

It should be noted when interpreting data from this study and others that the presence of macrophages within the cochlea does not correlate distinctly with an ongoing inflammatory response. Populations of native/resident macrophages have been found in different portions of the human cochleae in non-inflammatory conditions(Hough et al., 2022; Kaur et al., 2015; Liu et al., 2018; Okayasu et al., 2019). Additionally, macrophages may assume a multitude of phenotypes with varied pro- or anti-inflammatory roles and thus, the physiologic consequences of macrophage accumulation cannot exclusively be extrapolated as a maladaptive or pro-inflammatory and pro-fibrotic event(Wynn and Vannella, 2016). Following this, our data demonstrate that biomaterial surface types influence macrophage phenotype, altering cytokine production and fibroblast proliferation (pro-fibrotic phenotype).

Taken together, the results of this study support the contention that the intensity of the FBR is partially determined by the characteristics and composition of the implanted biomaterial. Fibrotic reactions are biomaterial specific, as demonstrated by the differences in cell adhesion, proliferation, and fibrosis on platinum versus PDMS. These data suggest that the inflammatory reaction to platinum contacts on CI electrodes likely contributes to fibrosis to a greater degree than previously thought. Further, the platinum contacts may also influence the deposition of new bone, as demonstrated in the *in vivo* data. This information can potentially be used to influence the design of future generations of neural prostheses. For example, thin film coatings of CI electrode arrays with ultra-low fouling biomaterials have the potential to dramatically reduce cellular adhesion to the electrode array surface(Leigh et al., 2017; Leigh et al., 2019; Shen et al., 2021). Another strategy to mitigate the cochlear tissue response to implanted electrode arrays is to delivery pharmaceuticals that modulate cellular responses. For example, electrode arrays that elute dexamethasone are currently under development in an effort to limit intracochlear tissue fibrosis(Farhadi et al., 2013; Toulemonde et al., 2021). Further studies are needed to explore strategies to limit the FBR to implanted biomaterials and improve hearing outcomes for CI recipients.

## Acknowledgements

We thank University of Iowa Small Animal Imaging Core for their expertise with 3D x-ray microscopy. This work was supported by grants #R01DC012578, R01DC018488, T32 DC00040 from the National Institutes of Health.

## References

1. Anderson, J.M., Rodriguez, A., Chang, D.T., 2008. Foreign body reaction to biomaterials. Semin Immunol 20(2), 86–100. https://doi.org/10.1016/j.smim.2007.11.004.

2. B. Majhy, P.P., A.K. Sen, 2021. Effect of surface energy and roughness on cell adhesion and growth - facile surface modification for enhanced cell culture. RSC Advances 11(25), 15467–15476.

3. Clark, G.M., Shute, S.A., Shepherd, R.K., Carter, T.D., 1995. Cochlear implantation: osteoneogenesis, electrode-tissue impedance, and residual hearing. Ann Otol Rhinol Laryngol Suppl 166, 40–42.

4. Claussen, A.D., Vielman Quevedo, R., Mostaert, B., Kirk, J.R., Dueck, W.F., Hansen, M.R., 2019. A mouse model of cochlear implantation with chronic electric stimulation. PLoS One 14(4), e0215407. https://doi.org/10.1371/journal.pone.0215407.

5. Deng, H., Maitra, U., Morris, M., Li, L., 2013. Molecular mechanism responsible for the priming of macrophage activation. J Biol Chem 288(6), 3897–3906. https://doi.org/10.1074/jbc.M112.424390.

6. Farhadi, M., Jalessi, M., Salehian, P., Ghavi, F.F., Emamjomeh, H., Mirzadeh, H., Imani, M., Jolly, C., 2013. Dexamethasone eluting cochlear implant: Histological study in animal model. Cochlear Implants Int 14(1), 45–50. https://doi.org/10.1179/1754762811Y.0000000024.

7. Foggia, M.J., Quevedo, R.V., Hansen, M.R., 2019. Intracochlear fibrosis and the foreign body response to cochlear implant biomaterials. Laryngoscope Investig Otolaryngol 4(6), 678–683. https://doi.org/10.1002/lio2.329.

8. Hallab, N.J., Bundy, K.J., O’Connor, K., Moses, R.L., Jacobs, J.J., 2001. Evaluation of metallic and polymeric biomaterial surface energy and surface roughness characteristics for directed cell adhesion. Tissue Eng 7(1), 55–71. https://doi.org/10.1089/107632700300003297.

9. Horzum, U., Ozdil, B., Pesen-Okvur, D., 2014. Step-by-step quantitative analysis of focal adhesions. MethodsX 1, 56–59. https://doi.org/10.1016/j.mex.2014.06.004.

10. Hough, K., Verschuur, C.A., Cunningham, C., Newman, T.A., 2022. Macrophages in the cochlea; an immunological link between risk factors and progressive hearing loss. Glia 70(2), 219–238. https://doi.org/10.1002/glia.24095.

11. Ishai, R., Herrmann, B.S., Nadol, J.B., Jr., Quesnel, A.M., 2017. The pattern and degree of capsular fibrous sheaths surrounding cochlear electrode arrays. Hear Res 348, 44–53. https://doi.org/10.1016/j.heares.2017.02.012.

12. Kamakura, T., Nadol, J.B., Jr., 2016. Correlation between word recognition score and intracochlear new bone and fibrous tissue after cochlear implantation in the human. Hear Res 339, 132–141. https://doi.org/10.1016/j.heares.2016.06.015.

13. Kan, C., Chen, L., Hu, Y., Ding, N., Lu, H., Li, Y., Kessler, J.A., Kan, L., 2018. Conserved signaling pathways underlying heterotopic ossification. Bone 109, 43–48. https://doi.org/10.1016/j.bone.2017.04.014.

14. Kaur, T., Zamani, D., Tong, L., Rubel, E.W., Ohlemiller, K.K., Hirose, K., Warchol, M.E., 2015. Fractalkine Signaling Regulates Macrophage Recruitment into the Cochlea and Promotes the Survival of Spiral Ganglion Neurons after Selective Hair Cell Lesion. J Neurosci 35(45), 15050–15061. https://doi.org/10.1523/JNEUROSCI.2325-15.2015.

15. Knoll, R.M., Trakimas, D.R., Wu, M.J., Lubner, R.J., Nadol, J.B., Jr., Ishiyama, A., Santos, F., Jung, D.H., Remenschneider, A.K., Kozin, E.D., 2022. Intracochlear New Fibro-Ossification and Neuronal Degeneration Following Cochlear Implant Electrode Translocation: Long-Term Histopathological Findings in Humans. Otol Neurotol 43(2), e153–e164. https://doi.org/10.1097/MAO.0000000000003402.

16. Leigh, B.L., Cheng, E., Xu, L., Andresen, C., Hansen, M.R., Guymon, C.A., 2017. Photopolymerizable Zwitterionic Polymer Patterns Control Cell Adhesion and Guide Neural Growth. Biomacromolecules 18(8), 2389–2401. https://doi.org/10.1021/acs.biomac.7b00579.

17. Leigh, B.L., Cheng, E., Xu, L., Derk, A., Hansen, M.R., Guymon, C.A., 2019. Antifouling Photograftable Zwitterionic Coatings on PDMS Substrates. Langmuir 35(5), 1100–1110. https://doi.org/10.1021/acs.langmuir.8b00838.

18. Li, P.M., Somdas, M.A., Eddington, D.K., Nadol, J.B., Jr., 2007. Analysis of intracochlear new bone and fibrous tissue formation in human subjects with cochlear implants. Ann Otol Rhinol Laryngol 116(10), 731–738. https://doi.org/10.1177/000348940711601004.

19. Liu, W., Molnar, M., Garnham, C., Benav, H., Rask-Andersen, H., 2018. Macrophages in the Human Cochlea: Saviors or Predators-A Study Using Super-Resolution Immunohistochemistry. Front Immunol 9, 223. https://doi.org/10.3389/fimmu.2018.00223.

20. Nadol, J.B., Jr., O’Malley, J.T., Burgess, B.J., Galler, D., 2014. Cellular immunologic responses to cochlear implantation in the human. Hear Res 318, 11–17. https://doi.org/10.1016/j.heares.2014.09.007.

21. Okayasu, T., O’Malley, J.T., Nadol, J.B., Jr., 2019. Density of Macrophages Immunostained With Anti-iba1 Antibody in the Vestibular Endorgans After Cochlear Implantation in the Human. Otol Neurotol 40(8), e774–e781. https://doi.org/10.1097/MAO.0000000000002313.

22. Okayasu, T., Quesnel, A.M., O’Malley, J.T., Kamakura, T., Nadol, J.B., Jr., 2020. The Distribution and Prevalence of Macrophages in the Cochlea Following Cochlear Implantation in the Human: An Immunohistochemical Study Using Anti-Iba1 Antibody. Otol Neurotol 41(3), e304–e316. https://doi.org/10.1097/MAO.0000000000002495.

23. Quesnel, A.M., Nakajima, H.H., Rosowski, J.J., Hansen, M.R., Gantz, B.J., Nadol, J.B., Jr., 2016. Delayed loss of hearing after hearing preservation cochlear implantation: Human temporal bone pathology and implications for etiology. Hear Res 333, 225–234. https://doi.org/10.1016/j.heares.2015.08.018.

24. Razafiarison, T., Holenstein, C.N., Stauber, T., Jovic, M., Vertudes, E., Loparic, M., Kawecki, M., Bernard, L., Silvan, U., Snedeker, J.G., 2018. Biomaterial surface energy-driven ligand assembly strongly regulates stem cell mechanosensitivity and fate on very soft substrates. Proc Natl Acad Sci U S A 115(18), 4631–4636. https://doi.org/10.1073/pnas.1704543115.

25. Scheperle, R.A., Tejani, V.D., Omtvedt, J.K., Brown, C.J., Abbas, P.J., Hansen, M.R., Gantz, B.J., Oleson, J.J., Ozanne, M.V., 2017. Delayed changes in auditory status in cochlear implant users with preserved acoustic hearing. Hear Res 350, 45–57. https://doi.org/10.1016/j.heares.2017.04.005.

26. Seyyedi, M., Nadol, J.B., Jr., 2014. Intracochlear inflammatory response to cochlear implant electrodes in humans. Otol Neurotol 35(9), 1545–1551. https://doi.org/10.1097/MAO.0000000000000540.

27. Shen, N., Cheng, E., Whitley, J.W., Horne, R.R., Leigh, B., Xu, L., Jones, B.D., Guymon, C.A., Hansen, M.R., 2021. Photograftable Zwitterionic Coatings Prevent Staphylococcus aureus and Staphylococcus epidermidis Adhesion to PDMS Surfaces. ACS Appl Bio Mater 4(2), 1283–1293. https://doi.org/10.1021/acsabm.0c01147.

28. Shepherd, R.K., Carter, P.M., Enke, Y.L., Thompson, A., Flynn, B., Trang, E.P., Dalrymple, A.N., Fallon, J.B., 2020. Chronic intracochlear electrical stimulation at high charge densities: reducing platinum dissolution. J Neural Eng 17(5), 056009. https://doi.org/10.1088/1741-2552/abb7a6.

29. Toulemonde, P., Risoud, M., Lemesre, P.E., Beck, C., Wattelet, J., Tardivel, M., Siepmann, J., Vincent, C., 2021. Evaluation of the Efficacy of Dexamethasone-Eluting Electrode Array on the Post-Implant Cochlear Fibrotic Reaction by Three-Dimensional Immunofluorescence Analysis in Mongolian Gerbil Cochlea. J Clin Med 10(15). https://doi.org/10.3390/jcm10153315.

30. Trouplin, V., Boucherit, N., Gorvel, L., Conti, F., Mottola, G., Ghigo, E., 2013. Bone marrow-derived macrophage production. J Vis Exp (81), e50966. https://doi.org/10.3791/50966.

31. Williams, D.F., 2009. On the nature of biomaterials. Biomaterials 30(30), 5897–5909. https://doi.org/10.1016/j.biomaterials.2009.07.027.

32. Witherel, C.E., Abebayehu, D., Barker, T.H., Spiller, K.L., 2019. Macrophage and Fibroblast Interactions in Biomaterial-Mediated Fibrosis. Adv Healthc Mater 8(4), e1801451. https://doi.org/10.1002/adhm.201801451.

33. Wynn, T.A., Vannella, K.M., 2016. Macrophages in Tissue Repair, Regeneration, and Fibrosis. Immunity 44(3), 450–462. https://doi.org/10.1016/j.immuni.2016.02.015.

